# Behavioural biomechanics: leaf-cutter ant cutting behaviour depends on leaf edge geometry

**DOI:** 10.1101/2024.12.06.626987

**Authors:** Frederik Püffel, Victor Kang, Mia Yap, Mohammad Shojaeifard, Mattia Bacca, David Labonte

## Abstract

Leaf-cutter ants cut fresh leaves to grow a symbiotic fungus as crop. During cutting, one mandible is typically anchored onto the leaf lamina while the other slices through it like a knife. When initiating cuts into the leaf edge, however, foragers sometimes deviate from this behaviour, and instead used their mandibles symmetrically, akin to scissors. *In-vivo* behavioural assays revealed that the preference for either of the two cutting strategies depended on leaf edge geometry, and differed between natural leaf margins that were straight or serrated with notch-like folds: leaf-cutter ants displayed a strong preference for scissor-cutting when leaf edges were straight or had wide notches. This preference, however, reversed in favour of knife-cutting when notches were narrow. To investigate whether this behavioural difference had a mechanical origin, we mimicked knife-cutting in *ex-vivo* cutting experiments: for wide notches, all but the sharpest mandibles failed to initiate cuts, or only did so at large forces, caused by substantial leaf buckling and bending. This increased force demand would substantially limit the ability of foragers to cut leaves, and so reduce the colony’s access to food sources. Scissor-cutting may thus be an adaptation to the mechanical difficulties associated with bending and buckling of thin leaves.

## 1 Introduction

Plant-feeding insects play a vital role in most terrestrial ecosystems: they are key pollinators of flowering plants [1], and they substantially accelerate the cycling of carbon, nitrogen and phosphor [2, 3]. However, they also destroy up to one-fifth of our global crop production [4–6], and so cause hundreds of US$ billions in annual economic loss [6, 7]. Key to their feeding success is the insects’ ability to overcome the plants’ mechanical defence mechanisms [8, 9]: slippery plant surfaces are adhered to via specialised tarsal structures [10, 11]; trichomes, small plant-epidermal extensions, are bypassed via elongated rostra [10, 12]; and wear-inducing leaf toughness is countered via incorporations of heavy metals, which harden the mandibular cutting edge [13–15]. This evolutionary ‘arms race’ is believed to have substantially driven the unparalleled diversification of both insects and plants [9, 16, 17].

Insect adaptations for plant-feeding are not only reflected by different genotypes; some developmental, morphological and behavioural adaptations occur plastically in direct response to varying mechanical demands and environmental constraints [18–22]: grass-feeding caterpillars and grasshoppers develop larger heads and mandible closer muscles when reared on ‘harder’ substrates [23, 24], allowing faster feeding rates [25]; butterfly larvae that feed on ‘tougher’ or silica-rich leaves grow to smaller sizes, but have longer and relatively heavier mandibles [26]; and cactus bugs grow mouthparts of varying length depending on the wall thickness of the fruits they feed on [27].

An insect herbivore that has adapted particularly well to the challenge of plant-foraging are leaf-cutter ants: they consume an approximate 15 % of the foliar biomass in the Neotropics [28]. Leaf-cutter ants cut small fragments from surrounding vegetation, carry them to a central nest, and incorporate them into a fungal garden grown as crop [29–31]. The intricate foraging ecology of leaf-cutter ants is characterised by an extensive division of labour and behavioural plasticity: larger ants cut and carry tougher leaves than smaller morphs from the same colony [29, 32, 33]; foragers cut smaller fragments when the food source is more attractive [34], closer to the nest [35], or when they face an uphill trail gradient [36] or height obstacles upon return [37]; longer fragments are carried at a steeper neck angle to maintain mechanical stability during walking [38, 39]; and foragers modify their leg extension to cut smaller fragments from thicker leaves [40].

The cutting of sheet-like vegetation gives rise to another mechanical challenge that a successful forager must overcome: leaf laminae are only a few hundred micrometers thick [41, 42], have weakly constrained margins, and thus bend and buckle readily upon the application of even minute forces [43]. How do leaf-cutter ants manage to avoid large deformations that would impede their ability to cut leaves?

Like most biting-chewing insects, leaf-cutter ants possess two mandibles that move predominately about a single joint axis of rotation [44–47]. The resulting planar kinematic space restricts the ant’s ability to manipulate food items [48], so that the propagation of cuts typically follows a simple stereotyped pattern: the ant anchors one of its mandibles onto the leaf lamina and pushes the other through the lamina akin to a knife slicing through a sheet of paper [49, 50]. When initiating cuts into straight leaf edges, however, we observed that ants often deviate from this ‘knife-cutting’ behaviour, and used both mandibles like a pair of scissors instead (figure 1b). After creating a small notch, the ant would then turn to stand on either side of the leaf lamina, and continue with knife-cutting. Experimental evidence suggests that introducing a small notch into the sheet edge prior to knife-cutting can mitigate out-of-plane deformations and facilitate cut initiation [50–53]. In addition to preliminary scissor-cuts, natural serrations of the leaf margin or previous cutting trajectories lead to variable leaf edge geometries that leaf-cutter ants may encounter during natural foraging. How does this variation, and the associated changes in mechanical constraints, affect the ants’ behaviour at cut initiation? Do leaf-cutter ants perhaps use scissor-cutting to avoid leaf bending and buckling?

**Figure 1.**
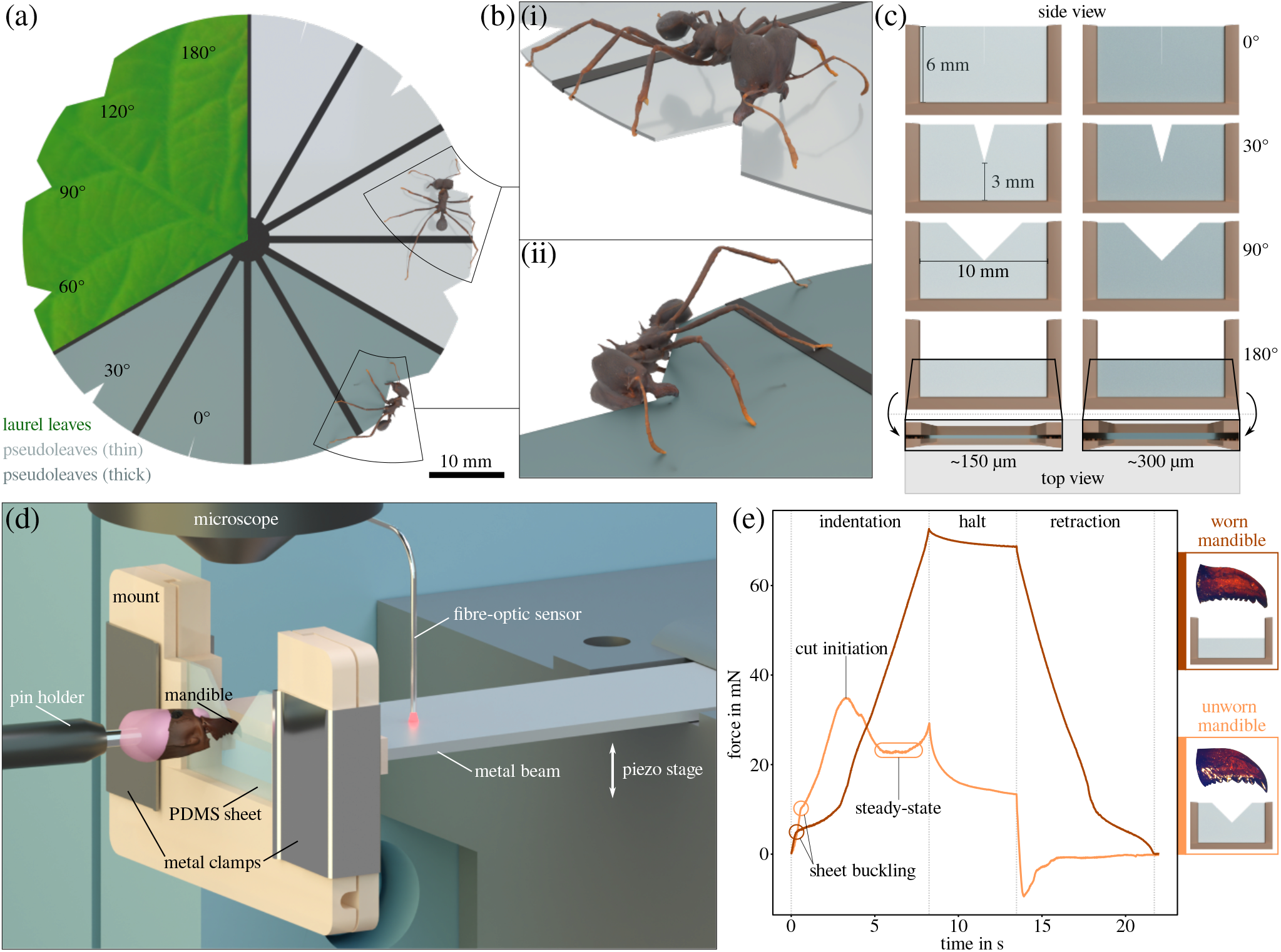
*(a)* To study the behavioural response of leaf-cutter ants to changes in the mechanical environment during leaf-cutting, we observed how *Atta cephalotes* workers initiated cuts at the edge of thin, leaf-like sheets with varying edge geometry. To this end, we prepared cutting discs (60 mm diameter) from laurel leaves and PDMS pseudoleaves, cut notch angles (0-180°) into the disc edges, and placed the discs inside the colony’s foraging arena. *(b)* From top-down video-recordings, two different initiation strategies were identified: (i) knife-cutting, where the long axis of the cutting mandible was perpendicular to the disc plane; and (ii) scissor-cutting, where the mandible long axes were approximately in parallel to the disc surface. The 3D reconstructions of the ant models used in this illustration were generated by Fabian Plum using *scAnt* [57]. *(c-d)* To investigate whether the choice of cutting strategy has a mechanical basis, we performed *ex-vivo* cut initiation force measurements using isolated mandibles (see [53] for further details on the sample preparation protocol and experimental setup). PDMS pseudoleaves of different thickness and notch angles were clamped and mounted onto the free end of a bending beam. The pseudoleaf was then moved against a stationary mandible via a piezo motor stage, so approximating the kinematics during knife-cutting. Upon contact between pseudoleaf and mandible, the beam deflected, as measured with a fibre-optic sensor. The deflection was then converted into a force using a calibration function. *(e)* Pseudoleaves were either cut (orange line representing an unworn mandible and a ∼150 *µ*m sheet with 90° notch angle), or merely deformed (mostly out-of-plane; brown line representing a worn mandible and a ∼150 *µ*m sheet with 180° notch angle); they often bent and buckled out-of-plane, visible by a sudden drop in ‘stiffness’ (circles). When a cut was initiated, the forces typically dropped to an approximately constant level after a peak at initiation (ellipse). When no cut was initiated, the forces instead continued to increase monotonously until the motor stopped; this typically happened when notch angles were large, pseudoleaves thin, or mandibles worn.

To address these questions, we investigated (i) whether the observed cutting behaviours occurred consistently across different substrates, (ii) how the behavioural preferences changed between serrated instead of straight-edged leaves, and (iii) how the presence and geometry of such serrations affected the biomechanics of cut initiation. By studying the cutting behaviour through a biomechanical lens, we aim to better understand the mechanisms that allow leaf-cutter ants to harvest plants effectively, and, more generally, how physical constraints can drive behavioural adaptations in insect herbivores.

## 2 Materials & methods

We initially studied the behaviour of leaf-cutter ants during cut initiation via *in-vivo* behavioural assays, where the colony was offered different substrates with varying notch geometries. Informed by observations from these assays, we then designed and performed *ex-vivo* cutting force experiments to rationalise the results, and to quantify mechanical difficulties at cut initiation.

### 2.1 Cutting substrate preparation

For behavioural experiments with freely foraging ants, we prepared cutting discs from spotted laurel leaves (*Aucuba japonica*), freshly collected from a single plant near campus (Imperial College London, South Kensington, UK). A circular paper template (60 mm diameter) was used to cut leaf discs from near the leaf base, and the lamina thickness was measured with a digital micrometre (0-25 mm range, ±1 *µ*m, Model 293-795-30, Mitutoyo Corporation, Kawasaki, Japan). The average leaf thickness was 390 ± 58 *µ*m (n=52 leaf discs; four replicates per leaf).

During natural foraging, leaf-cutter ants always initiate cuts from the leaf edge [54, 55]. The geometry of this edge, however, can vary: for entire leaves with intact margins, the leaf edges are approximately straight; for leaves with serrated margins, at the intersections of previous cutting trajectories, or when a scissor-cut was made first, the leaf edge has notch-like folds instead. To systematically control for the variation of leaf edge geometry, we cut notches of consistent depth (3 mm), comparable to the mandible length of large *Atta* foragers [∼2 mm, 56] into each of twelve disc sections using a scalpel blade (carbon steel, no. 11, Swann-Morton, Sheffield, UK). Notch angles ranged from a single slit (0° notch angle) to an intact straight edge (180° notch angle — i. e. no notch), with intermediate notch angles of 30°, 60°, 90°, and 120°, respectively (figure 1a).

In addition to experiments with laurel leaves, we also used thin sheets of a silicone polymer as cutting substrates (poly-dimethylsiloxane, PDMS). Synthetic ‘pseudoleaves’ were developed in an effort to minimise uncontrolled co-variation present in biological materials: pseudoleaves are homogeneous and can be fabricated with consistent thickness; moreover, PDMS has well-defined mechanical properties, enabling theoretical predictions on cutting force and buckling behaviour. When scented, pseudoleaves are also readily cut by leaf-cutter ant foragers, making them a powerful tool for behavioural assays [35, 40].

PDMS sheets were made with a 10:1 mixing ratio (base:curing agent; Sylgard 184, Dow Inc., MI, USA), degassed in a vacuum chamber and sandwiched between two silanised glass plates, separated by feeler gauges (150, 200, 300 or 500 *µ*m; Precision Brand, Downers Grove, IL, USA; see [53]). All PDMS samples were cured in an oven at 65°C for four hours. For *in-vivo* behavioural assays, a 3D-printed support structure was embedded into the PDMS prior to curing. This structure consisted of 12 radial spokes, which spanned across a diameter of 60 mm, so creating wheel-like cutting discs akin to leaf laminas supported by veins (figure 1a). We cast cutting discs of two different thicknesses: 160*±*4 *µ*m (mean*±*standard deviation) and 310*±*5 *µ*m, respectively (n=3 pseudoleaf discs per thickness). Sheet thickness, *t*, strongly influences its flexural rigidity [*D* ∝ *t*^3^, 43], and thus the resistance against out-of-plane deformation. In analogy to the cutting discs made from laurel, notches were cut into the edges of the cured pseudoleaves. Prior to the behavioural experiments, the pseudoleaf cutting discs were cleaned from any contaminants using 50 % ethanol, rinsed with purified water, immersed in honey water for at least 5 min, and left to dry at ambient conditions.

For *ex-vivo* cut initiation experiments (see below), PDMS sheets of similar thickness were cast (156*±*3 *µ*m and 305*±*3 *µ*m, respectively, n = 72 for each), and cut with a scalpel blade to rectangles of 20 × 9 mm (width × height), with central notches of 3 mm length and 0°, 30°, 90°, or 180° notch angles. To relate cutting forces to the mechanical properties of PDMS, pure shear tests were performed following the protocol described in Püffel et al. [53]. The average critical strain energy release rate, or fracture toughness, was 156*±*27 J/m^2^ (n = 6, see [52]; the average thickness of the pure shear test samples was 508*±*7 *µ*m).

### 2.2 Study animals

A mature *Atta cephalotes* colony was used for all experiments. The colony was kept in a climate chamber at 25°C and 60% relative humidity (FitoClima 12.000 PH, Aralab, Rio de Mouro, Portugal), and fed with bramble, laurel leaves, and maize *ad libitum*.

### 2.3 Cutting-behavioural assays

*In-vivo* cutting assays were conducted in a foraging arena (30 × 22 cm), connected to the nest via ≈5 m of transparent plastic tubing (24 mm inner diameter). A video camera was positioned above the arena (HQ camera controlled via Raspberry Pi 3A+; Raspberry Pi Foundation, Cambridge, UK), and the camera field of view, focus and exposure times were adjusted prior to each trial. Cutting discs were placed inside the arena on a small 3D-printed ‘stool’, to prevent contact between disc and arena floor. Videos were recorded at 10 fps until cuts had been initiated in each of the twelve disc sections. For each section, the mode of cut initiation was extracted, based on the orientation of the mandible long axis relative to the leaf plane (see below). For all notch angles but 180° (straight edge), only cuts made at the notch centre were considered for further analysis to minimise confounding effects due to variation in boundary conditions. Experiments were terminated after at least ten valid cuts per notch angle and substrate were recorded. The median number of valid trials across test conditions was 42, with a minimum and maximum of 10 and 47, respectively.

Per our preliminary observations, ants initiated cuts using one of two strategies (figure 1b): (i) the ant stood on either side of the disc with the long axes of its mandibles approximately perpendicular to the substrate plane. One mandible served as a fixed anchor point towards which the other mandible was then moved to cut the leaf, starting from the leaf edge (knife-cutting, see above and [49, 50]). Or (ii), the ant stood on the disc edge with the long axes of its mandibles orientated in parallel to the disc plane, and cuts were made by drawing the mandibles together symmetrically (scissor-cutting). The choice of cut initiation strategy was easily identifiable from the videos.

### 2.4 Cut initiation force experiments

To probe for a potential mechanical basis for a preference in cut initiation strategy, we performed *ex-vivo* cut initiation force measurements with isolated mandibles. The goal was to create a cutting scenario that was mechanically similar to knife-cutting. To this end, we used an experimental setup initially developed to measure steady-state cutting forces (figure 1d and [53]). In brief, the setup consisted of a fibre optic displacement sensor that measured the deflection of a bending beam (*µ*DMS-RC32 controlled via DMS Control v 3.015, Philtec, Annapolis, MD, USA; see electronic supplementary material for details on sensor calibration and drift correction). Attached to this beam was a polymer mount that held a PDMS pseudoleaf via two metal clamps. Pseudoleaf, beam, and sensor were moved upwards by a piezo motor, such that the pseudoleaf was pressed against a stationary mandible, causing the beam to deflect. The mandible was still attached the head capsule, which was glued onto an insect pin connected to a 3D micro manipulator (see below for details on mandible preparation). The mandible was positioned such that its cutting edge was perpendicular to the sheet, with its distal-most part slightly extending over the sheet edge (akin to knife-cutting, figure 1d). The polymer mount, which held the PDMS pseudoleaf, was U-shaped, with a ‘free’ cutting region of 10 × 6 mm (width × length), large in comparison to the sheet thickness (150-300 *µ*m), and similar to the circumferential distance between neighbouring spokes in the cutting discs (*π*60 mm*/*12 ≈ 16 mm). The PDMS sheets were clamped such that the notch was in the centre of the mount, 3 mm away from the lower clamping bar (figure 1c). This distance was chosen to be approximately equal to the maximum mandibular gape of a large *Atta* forager [58]; the clamping bar thus served a similar mechanical function to the anchoring mandible during cut initiation. Experiments that simulate scissor-cutting were attempted, but abandoned due to substantial technical difficulties associated with small ant heads and brittle apodemes [59].

A total of 18 mandibles were used for these experiments, collected from 18 ant workers that were extracted from a foraging arena connected to the nest by about ≈0.5 m of tubing (the arena contained small amounts of fungus, so that some of the collected ants may not have been actively foraging). Ants were sacrificed by freezing, weighed to the nearest 0.1 mg (Explorer Analytical EX124, max. 120 g × 0.1 mg, OHAUS Corporation, Parsippany, NJ, USA; body masses ranged between 17.0 mg and 37.3mg), and dissected following the protocol described in Püffel et al. [53]. Prior to the cut initiation experiments, individual mandibles were photographed, and steady-state cutting forces measured using PDMS pseudoleaves of an intermediate thickness (207±3 *µ*m). Depending on the extent of mandibular wear, steady-state cutting forces can vary by a factor of about seven among similarly-sized *Atta* ants [50, 53]. Extracting steady-state in addition to initiation forces thus allowed us to (i) determine the relative force increase at cut initiation compared to steady-state (see below for details on the calculation), (ii) link the effects of mandibular wear to the probability of pseudoleaf buckling and cut initiation, and (iii) speculate on how mechanics may affect ontogenetic changes in the ants’ cutting behaviour. These separate experiments were necessary, as it was generally not possible to directly measure steady-state cutting forces from the cut initiation experiments; cuts were often not initiated at all, or if they were, the steady-state region was too short to confidently extract an average force (see electronic supplementary material for more details).

Cut initiation force experiments involved eight measurements per mandible, one for each notch angle and sheet thickness, in randomised order. Individual mandibles were mounted, oriented using a top-down microscope, and positioned such that the cutting edge was close to the notch centre (<50*µ*m distance, controlled via the motor stage). Subsequently, the force recording was started, and the beam mount was moved against the mandible at a constant rate of 0.3 mm/s, corresponding to the upper end of observed leaf-cutting speeds for larger foragers [32, 60]. The motor moved a total distance of 2.5 mm (‘indentation’ phase in figure 1e). To avoid potential damage to the mandible, the motor stopped at a distance of ≈0.5 mm to the lower clamping bar (‘halt’ phase). The motor stage then returned to its original position (‘retraction’ phase), and the PDMS sheet was inspected through a microscope to determine if a cut was initiated. Measurements were retaken if: (i) the head capsule came into contact with the mount prior to cut initiation; (ii) the mandible-pin complex moved out of the pin holder; or, (iii) the mandible slipped out of the notch, which typically occurred after substantial pseudoleaf buckling.

Where cuts were successfully initiated, we extracted the cut initiation force, *F*_*i*_, from the drift-corrected force-time graph. *F*_*i*_ was identified by a sharp drop in ‘stiffness’, followed by a short, approximately steady-state region (figure 1e and [61–63]); *F*_*i*_ typically corresponded to the maximum measured force during the experiment. The average cutting force was extracted from the steady-state force experiments, and rescaled in proportion to sample thickness (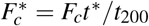— cutting forces increase in direct proportion to sheet thickness, see [64], and Walthaus et al., in preparation, for experimental verification using *Atta* mandibles). The relative force increase was then calculated as 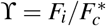. Normalising cut initiation forces with steady-state forces allowed us to account for inter-individual differences in mandibular wear; a worn mandible requires higher forces to initiate a cut than an unworn mandible, but it also propagates cuts with larger steady-state forces.

### 2.5 Finite Element Analysis

Key to cutting success is to avoid large out-of-plane deformations during cut initiation. We systematically tested how notch angle and sheet thickness influences the leaf’s resistance against such deformations by performing Finite Element Analysis: 3D models of both radial disc sections, used for *in-vivo* trails, and rectangular sheets, used for *ex-vivo* cutting experiments, were constructed in Abaqus/CAE (SIMULIA, Dassault Systèmes SE, Vélizy-Villacoublay, France). Suitable mesh densities were determined via preliminary convergence study; the resulting meshes, for which further refinement was ineffectual, consisted of ∼136,000 C3D8RH elements (∼152,000 nodes) for radial models, and ∼173,000 C3D8RH elements (∼194,000 nodes) for rectangular models. Across both geometries, two different sheet thicknesses (150 and 300 *µ*m) and seven notch angles (0°, 30°, 60°, 90°, 120°, 150°, and 180°) were simulated, leading to a total of 28 simulations. The material was modelled as linear elastic, and simulations were completed using the Static General step. The experimental clamping conditions were mimicked by fixing the node displacements along the left, right, and bottom edges in all six degrees of freedom (see figure 1a,c). For each simulation, a uniform out-of-plane displacement (200 *µ*m) was applied to the notch centre, and the resulting reaction forces were extracted. Out-of-plane stiffness was calculated as out-of-plane reaction force divided by the imposed displacement.

In addition to *out-of-plane* deformations, cut initiation also depends on the sheet’s *in-plane* stiffness and the concentration of tensile stresses for crack nucleation [65, 66]. We quantified the effects of notch angle and sheet thickness on in-plane stiffness, and the resulting tensile stresses, by performing a second set of simulations on all rectangular models (14 simulations). The simulation parameters were equal to those used for out-of-plane simulations, except that the displacement at the notch centre (200 *µ*m) was applied in the cutting direction (in-plane). We calculated the in-plane stiffness as in-plane reaction force divided by the imposed displacement, and extracted the maximum tensile stress in the pseudoleaf from each simulation.

### 2.6 Statistical analysis

The relationship between leaf edge geometry and cut initiation strategy was quantified with a binary logistic regression analysis, using notch angle and sheet thickness as independent variables. The logistic function took the form *P*(*Y*) = 1*/*[1 + *exp*(−(*b*_0_ + *b*_1_*X*_1_ + … + *b*_*n*_*X*_*n*_))], where *P*(*Y*) is the probability of a knife-like cut initiation, *X*_*i*_ represent the predictors (notch angle in degrees and sheet thickness in micrometers), and *b*_*i*_ are the respective regression coefficients. Results from *ex-vivo* cutting experiments were analysed with repeated measures binary logistic regression analysis, again using notch angle and sheet thickness as fixed effects, and mandible as random effect. Here, the dependent variable encoded whether or not a cut was initiated. To quantify the effects of sheet geometry on the mechanical difficulty to initiate cuts, a linear mixed model was used only on those data where cuts were successfully initiated; the model used the relative force increase, ϒ, as dependent variable, notch angle and sheet thickness as fixed effects, and mandible as random effect. Model goodness of fit was assessed via a *χ*^2^-test and the Akaike Information Criterion (AIC). All statistical analyses were conducted in R v4.1.2 [67], following Field et al. [68]; the R package *lme4* v1.1-31 was used for mixed models.

## 3 Results

### 3.1 Leaf-cutter ants systematically prefer scissor-cuts for sheets with wide notches

*In-vivo* cutting behaviour experiments revealed that leaf-cutter ants virtually always used scissor-cutting to initiate cuts into straight leaf edges for both real and pseudoleaves (180° notch angle, figure 2a). This strong preference steadily declined with decreasing notch angles, and ultimately reversed in favour of knife-cutting for 0° notch angles, where only about 10 % of the workers used scissor-cutting to initiate cuts. Accordingly, the logistic regression coefficients for notch angle were significantly negative for both pseudo- and laurel leaves (*b*_1_ = -0.0301 (standard error, SE: 0.0030) and *b*_1_ = -0.0277 (SE: 0.0041), respectively; see electronic supplementary material table S1 for detailed statistics) — the wider the notch angle, the lower the probability of a knife-cut. The notch angles at which both cut initiation strategies occurred with equal probability, *P*(knife-cut) = *P*(scissor-cut) = 0.5, were 58° (thin pseudoleaves), 38° (thick pseudoleaves), and 39° (laurel leaves), respectively. The preferred strategy also depended significantly on pseudoleaf thickness (*b*_2_ = -0.0040 (SE: 0.0015)): cuts into thick pseudoleaves were slightly less likely to be initiated by a knife-cut than cuts into thin pseudoleaves. However, the associated change in odds per unit increase in thickness, *P*(knife-cut)*/P*(scissor-cut), was small (0.996 (confidence interval, CI: 0.993|0.999), where a value of one indicates no change).

**Figure 2.**
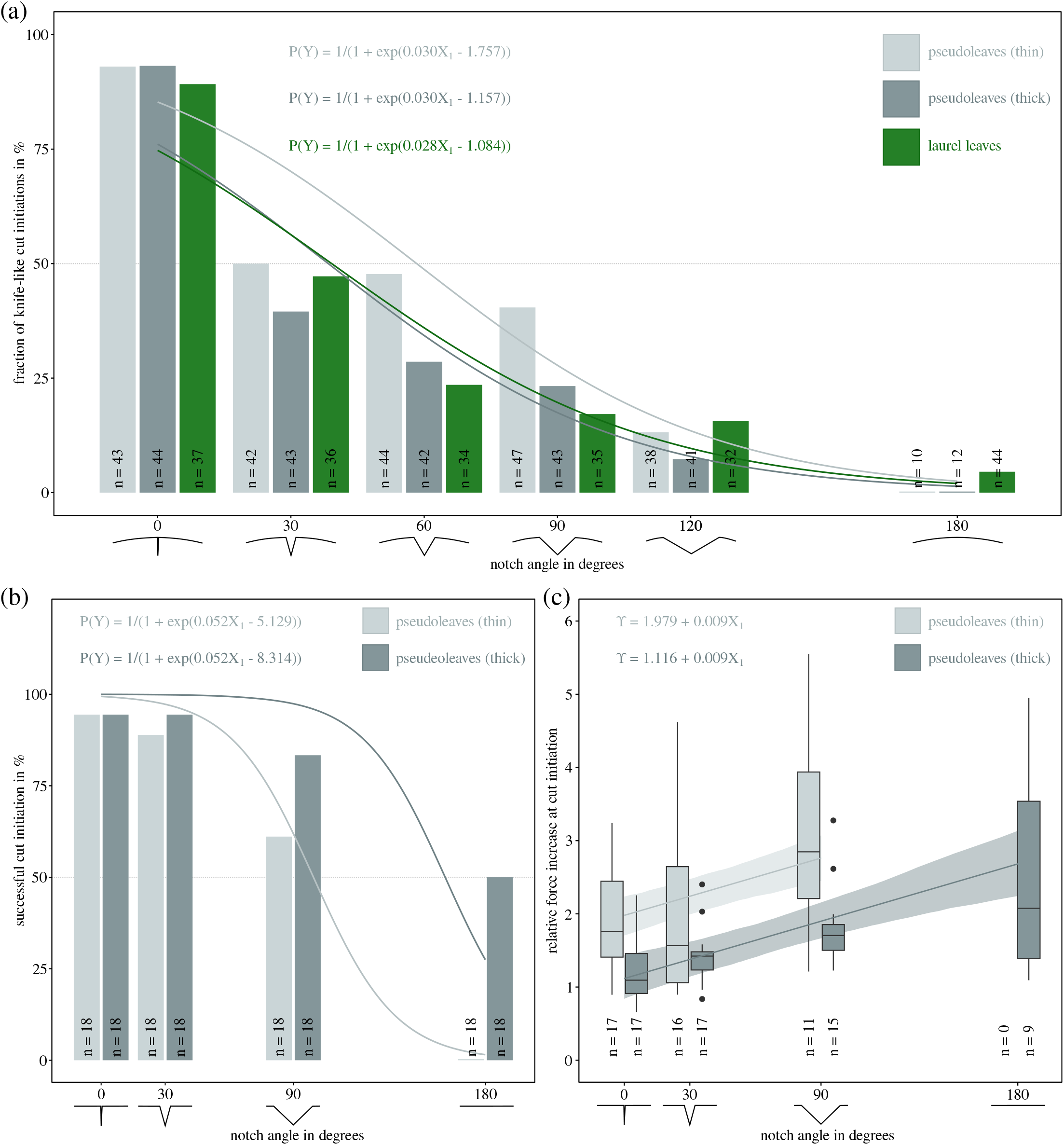
*(a)* Leaf-cutter ants prefer to initiate cuts with scissor-cutting when notch angles are large, but with knife-cutting when notch angles are small. A binary logistic regression analysis revealed a significant decrease in probability of knife-cut initiation (*P*(*Y*)) with increasing notch angles (*X*_1_; *b*_1_ = -0.0301 (standard error, SE: 0.0030) for PDMS pseudoleaves, and *b*_1_ = -0.0277 (SE: 0.0041) for laurel leaves). Experiments with PDMS pseudoleaves showed that *P*(*Y*) also varied significantly with pseudoleaf thickness, but this effect was small (*X*_2_; *b*_2_ = -0.0040 (SE: 0.0015)). *(b)* The change in cut initiation preference with edge geometry can be rationalised via results obtained from *ex-vivo* cut initiation experiments, designed to imitate knife-cutting. Successfully initiating a cut via knife-cutting was significantly less likely for larger notch angles (*P*(*Y*); *b*_1_ = -0.0515 (SE: 0.0129)), and significantly more likely for thicker pseudoleaves. Thin pseudoleaves instead often deformed substantially out-of-plane (see supplementary figure S1; *b*_2_ = 0.0214 (SE: 0.0067))). *(c)* Even where cuts were initiated, this involved substantially larger forces, as confirmed by the ratio between cut initiation and steady-state cutting forces, ϒ. ϒ increased significantly with notch angle (0.0087 (SE: 0.0017) per degree angle change), and decreased significantly with sheet thickness (−0.0058 (SE: 0.0012) per micrometer thickness change). The increase in cut initiation force may deter if not physically prevent leaf-cutter ants from using knife-cutting to initiate cuts into straight leaf edges. This limitation may be particularly restrictive for small ants with worn mandibles, which require a larger fraction of their total bite force capacity to cut the same material [53, 58].

### 3.2 Buckling impedes knife-like cut initiation for wide notches

To probe for a potential mechanical underpinning of the behavioural preference for scissor-cuts for large notch angles, we conducted *ex-vivo* cutting force experiments that imitated knife-cutting (figure 1d). When pseudoleaves had wider notches, cuts were less likely to be initiated (figure 2b): a unit increase in notch angle was associated with a significant decrease in cut initiation probability (*b*_1_ = -0.0515 (SE: 0.0129)). Sheet thickness, in turn, had a significant positive effect on initiation probability (*b*_2_ = 0.0214 (SE: 0.0067)): the thicker the sheet, the higher the probability of cut initiation. The notch angles at which cuts were initiated half of the time were 100° and 161°, for thin (∼150 *µ*m) and thick (∼300 *µ*m) pseudoleaves, respectively. Where cuts failed to initiate, the mandible remained in contact with the notch centre throughout the experiment, but the pseudoleaf deformed substantially out-of-plane instead of being cut (see supplementary figure S1 and 1e for a typical force profile). Once the motor had returned to its original position, the intact pseudoleaf flattened again, indicating that its deformation had been predominantly elastic. Out-of-plane deformation occurred often even when cuts were successfully initiated. Cut initiation was then typically associated with a sudden reduction in out-of-plane deformation. In cases where some out-of-plane deformation remained visible at the end of the indentation phase, the associated cuts tended to be shorter, as assessed via visual inspection. Shorter cuts and a delayed cut initiation typically occurred for mandibles that required larger steady-state cutting forces (figure 3).

**Figure 3.**
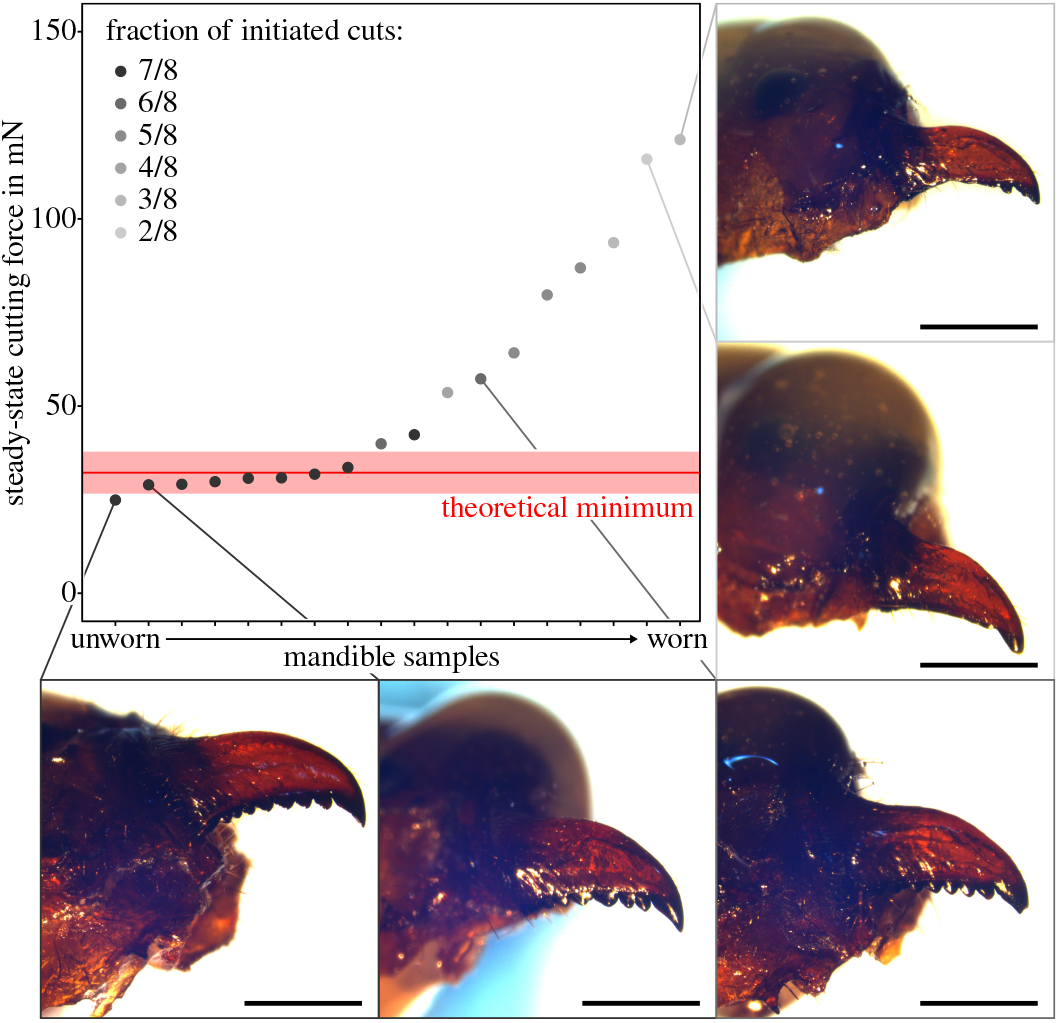
To obtain a proxy for mandible sharpness, steady-state cutting forces were measured for each mandible. Across all mandibles, steady-state forces ranged between 25 and 121 mN. Almost half of the mandibles cut with approximately the same force, close to a theoretical minimum predicted from the PDMS fracture toughness and sheet thickness (32±6 mN; the shaded area represents the standard deviation calculated as propagated uncertainty following [69]); these mandibles may be considered ‘ideally sharp’ [53]. Larger steady-state forces accordingly occurred for mandibles that were visibly worn; such mandibles also had a significantly lower probability to initiate cuts across all test conditions. Mandibles that required the highest steady-state forces had ‘lost’ almost all their teeth apart from the (distal-most) apical tooth. Scale bars represent a length of 1 mm.

### 3.3 Knife-like cut initiation requires higher forces when notches are wider

To quantify the mechanical difficulty of cut initiation, we calculated the relative force at initiation compared to steady-state cutting. Steady-state cutting forces ranged from 25 to 121 mN, mirroring previous results [53, and figure 3]. The smallest cutting forces were close to a theoretical minimum estimated from a simple mechanical model that uses PDMS fracture toughness *G*_*c*_ and pseudoleaf thickness *t* [53, 64], to estimate *F*_*min*_ = *G*_*c*_*t*_200_ = 32±6 mN (the standard deviation was calculated as propagated uncertainty following [69]). Mandibles with high steady-state cutting forces had worn and sometimes completely ablated teeth (figure 3), and initiated significantly fewer cuts (Kendall rank correlation: *τ* = −0.76, p < 0.0001, n = 18). Across all successful cuts, the relative force increase was 0.7 ≤ ϒ ≤ 5.6 (in 13 out of 102 measurements, *F*_*i*_ was slightly lower than the steady-state force, 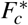 in all other cases, ϒ *>* 1). The smallest force increase was observed for thick pseudoleaves with narrow notches, where ϒ ≈ 1 — the force at cut initiation was about equal to the steady-state cutting force. ϒ increased significantly with notch angle (0.0087 (SE: 0.0017) per degree change in angle), approaching 3 for thick pseudoleaves with straight edges (figure 2c): the wider the notch, the more adverse the effect of sheet deformation on cut initiation force. ϒ was also higher for thinner pseudoleaves: cut initiation forces were approximately double the steady-state cutting force for thin pseudoleaves with 0° notch angles (ϒ ≈ 2), and increased to almost three times compared to the steady-state cutting force for 90° notch angles (slope: -0.0058 (SE: 0.0012) per micrometer change in thickness). ϒ could not be evaluated for thin pseudoleaves with straight edges, because none of the 18 tested mandibles successfully initiated a cut. Both notch angle and sheet thickness thus had a significant effect on ϒ; accordingly, the variation of ϒ was best described by a linear mixed model with both notch angle and sheet thickness as fixed effects, *f* (*X*_1_, *X*_2_), compared to a model with notch angle as single predictor, *f* (*X*_1_), or an intercept-only model (*f* (*X*_1_) vs ‘intercept’: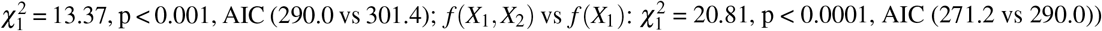.

### 3.4 Out-of-plane and in-plane stiffness decrease substantially with increasing notch angle

The effects of notch angle on the sheet’s resistance against and out-of-plane and in-plane deformation was further investigated with Finite Element Analysis. Out-of-plane stiffness varied strongly between thin and thick pseudoleaves across both geometries: for radial disc sections, it was 2.68±0.02 times larger for thicker sheets, almost identical to the results obtained for rectangular sheets (2.66±0.06; see figure 4). The out-of-plane stiffness was also substantially affected by notch angle: across both geometries and sheet thicknesses, out-of-plane stiffness was maximum for sheets with sharp notches, and monotonously decreased with increasing notch angle to less than one-third for straight leaf edges (≈0.31 and ≈ 0.17 for radial and rectangular geometries, respectively). The largest drop in stiffness between two consecutive angle increments occurred consistently between 150° and 180°, with a drop of ≈55% and ≈71% for radial and rectangular geometries, respectively.

**Figure 4.**
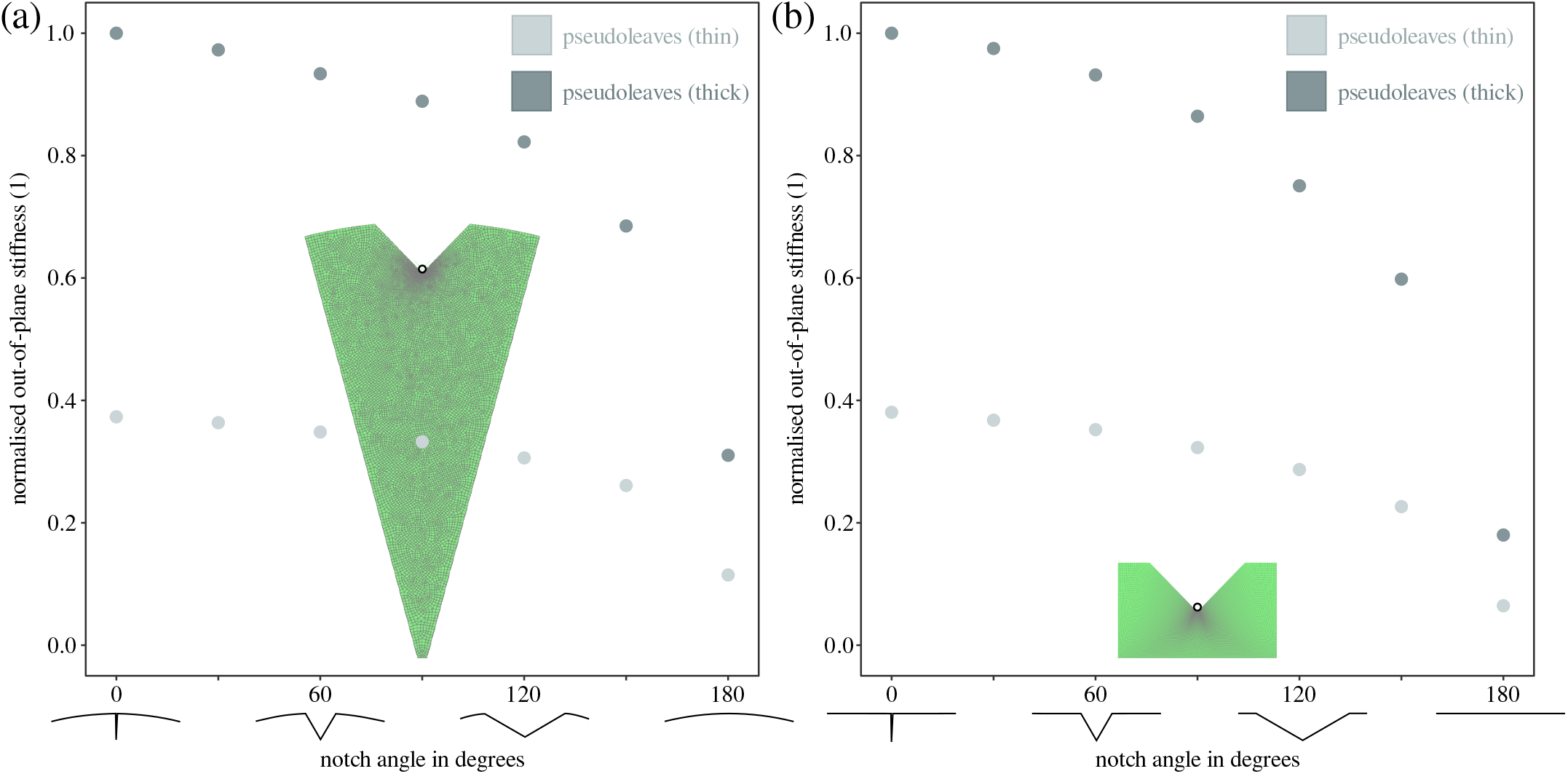
To further explore the mechanical underpinning of the observed differences in cutting strategy and cut initiation forces, we performed Finite Element Analysis. 3D models of both radial disc sections— *(a) in-vivo* behavioural assays— and rectangular sheets—*(b) ex-vivo* cutting experiments— were subjected to a fixed out-of-plane displacement at the notch centres. The resulting reaction force per unit displacement, the out-of-plane stiffness, was maximum for thick sheets with narrow notches, and monotonously decreased with increasing notch angle. For both radial and rectangular geometries, the values for out-of-plane stiffness were normalised with their respective maxima. In general, thick sheets had a larger out-of-plane stiffness than thin sheets (≈2.7 times). Across both geometries and sheet thicknesses, the out-of-plane stiffness decreased by a factor 3-6 between 0° and 180° notch angles, suggesting that sheets with narrow notches have a substantially higher resistance against bending and buckling. This increased resistance against out-of-plane deformation, in conjunction with higher maximum tensile stresses (see electronic supplementary materials figure S2b), likely facilitates knife-like cut initiation, in line with both behavioural observations— knife-cuts were preferred for sheets with sharp notches (figure 2a), and cutting experiments— the force at cut initiation was lowest for 0° notch angles (figure 2c).

In analogy to these results, sheet thickness and notch angle also affected the resistance against in-plane deformation. In-plane stiffness was consistently larger for thicker sheets (1.99*±*0.03; see electronic supplementary material figure S2a), and monotonously decreased from its maximum at 0° notch angle to approximately half of that for straight edges. As a result of this increased in-plane stiffness, the maximum tensile stresses were substantially larger for sharp notches than for wide notches at equivalent indentation depths (factor of ≈3 between 0° and 180° notch angles, see electronic supplementary material figure S2b).

## 4 Discussion

The mechanical demands of plant-foraging have shaped the evolution of mouthpart morphology and the feeding behaviour across the herbivorous Insecta [10, 46, 48, 70, 71]. Some herbivores, such as leaf-cutter ants, display a diverse set of adaptations to meet these demands effectively: leaf-cutter ant colonies deploy a polymorphous workforce to forage on plant materials of variable properties, where larger workers cut and carry tougher leaves [29, 32, 33]; the musculoskeletal bite apparatus is specialised to produce exceptionally large weight-specific bite forces [56, 58, 72, 73], and to cut leaves with lower forces than pristine scalpel blades [50, 53]; leaf-cutter ant foragers favour pre-cut leaf fragments over intact leaves which would require more cutting [74]; and they adjust their gaits and neck positions to the size and shape of the carried fragments [39].

In this study, we show that leaf-cutter ants also adjust their cutting strategy to overcome a mechanical difficulty specific to thin-sheet cutting: workers tend to prefer scissor-over knife-cuts when the risk of out-of-plane deformation is high. This strong behavioural preference can be rationalised with results from *ex-vivo* cutting experiments that mirrored knife-cutting: an increase in notch angle was associated with a decrease in initiation probability, and a substantial increase in the cut initiation force. In the following paragraphs, we discuss the adverse consequences of increased cut initiation forces for the colony’s ability to access food sources, and explore the potential mechanical basis for behavioural plasticity during cut initiation.

### 4.1 Lower cut initiation forces may increase the range of accessible food sources

Leaf-cutter ants can only cut leaves if they can generate sufficiently large bite force. The required bite force generally depends on the material properties of the leaf, and the geometry of the cutting mandible. Based on *in-vivo* bite force measurements [58], combined with leaf-mechanical data [41], we previously estimated the typical force a leaf-cutter ant forager may have to generate to cut the median tropical leaf as about 82 mN [58]. The force at cut initiation for a 0° notch was ϒ = 2.883 + 0.009(0°)−0.006(median leaf thickness = 210 *µ*m) ≈1.67 larger than the typical steady-state force measured for the same material. Extrapolating from this data, the minimum ant body mass necessary to initiate a cut via knife-cutting may be estimated from the bite force data as about 6 mg. For a straight leaf edge (180° notch angle), this force would increase instead by a factor ϒ≈ 3.24 and the minimum body mass to about 12 mg — twice as heavy as for a narrow notch, and almost four times as heavy as required for steady-state cutting forces. If these differences appear small, consider the reduction in accessible plant material: a worker ant with a body mass of 6 mg can generate bite forces sufficient for the steady-state cutting of 78 % of tropical leaves, but can only initiate knife-cuts into a meagre 16 % — and no steady-state cutting is possible without initiating a cut. The large cut initiation forces associated with out-of-plane sheet deformation may thus effectively prevent small ants from cutting leaves with straight edges altogether. Larger ants are likely less affected by this force increase: due to an increased bite force capacity, a 30 mg ant could still initiate knife-cuts into 88 % of tropical leaves with straight edges, compared to 98 % for leaves with narrow notches. However, the increased forces and associated energetic costs may still drive large ants to use scissor-cutting instead; we caution against definitive conclusions on the energetic benefits of scissor-cutting, which require either *in-vivo* metabolic rate measurements during cut initiation, or experiments with a force measurement setup that imitates scissor-cutting (see discussion below).

An ant’s ability to initiate knife-cuts is affected not only by its body size, but also by the wear state of its mandibles. Sharp mandibles, with low steady-state cutting forces, successfully initiated cuts across all test conditions, except for thin sheets with straight edges. In contrast, worn mandibles with high steady-state forces only initiated cuts when notches were narrow (≤ 30°, figures 2b and 3). Knife-like cut initiation is thus particularly difficult for foragers with worn mandibles, and this difficulty may drive older ants to avoid this strategy altogether [50, 75, 76]. Across active foragers, the distribution of mandibular wear states may be partially reflected by the statistical distribution of behavioural responses observed during the cutting assays (figure 2a): ants with particularly worn mandibles may comprise the small fraction of foragers that still deployed scissor-cutting even when notch angles were small. Conversely, the few ants that used knife-cutting for pseudoleaves with wide notches may have possessed almost pristine mandibles, so that cut initiation occurred at loads too small to cause excessive out-of-plane deformations. In addition to variation in mandibular wear, the ants’ behavioural preference may also be influenced by size-related differences in bite force (see above), individual preferences, and local environmental conditions: narrow notches may render the ant’s access to the notch centre difficult, so that knife-cuts are simply more practical. The ability to anchor the legs onto the sheet edge, as well as interactions with other ants may have further contributed to a ‘softer’ transition between the two initiation strategies.

### 4.2 Biomechanics of cut initiation - what are the advantages of scissor-cuts?

The results from *ex-vivo* cutting experiments provide strong arguments *against* the use of knife-cutting when leaf edges are straight or have wide notches: large out-of-plane deformations lead to increased cut initiation forces or even prevent cut initiation altogether. But what are the arguments *for* using scissor-cutting instead?

To approach this question from a mechanical angle, we first note that knife- and scissor-cutting are not mechanically equivalent— they differ in their respective modes of fracture (mode I vs mode III), blade orientation relative to the sheet plane, and ‘slice-to-push’ ratios [66, 70, 77–79]. However, both cutting modes involve material fracture and thus the creation of new surface area. Each unit area, *dA* of new surface demands a minimum energy investment, *dU*_*A*_ = *G*_*c*_*dA*, which must be supplied by the cutting mandible; additional energy sinks may emerge from friction and non-recoverable elastic strain energy [64, 77]. Although simple, such virtual work arguments have been successfully used to demonstrate that pristine leaf-cutter ant mandibles are close to optimally ‘sharp’, i. e. steady-state forces during knife-cutting of PDMS pseudoleaves were close to a theoretical minimum, *F*_*c*_ ≈ *F*_*min*_ = *G*_*c*_*t* [53].

During scissor-cutting, additional energetic costs may arise from friction between the overlapping blades: industrial scissors are often spring-loaded to enable ‘clean’ cuts of thin sheets [79–81], and leaf-cutter ant mandibles contact and elastically pitch out-of-plane when they overlap [47]. Even if this frictional contribution was negligible, the energy required to scissor-cut would still be bound from below by *G*_*c*_*dA*, the minimum achieved by almost half of the mandibles during knife-cutting (figure 3). For sharp mandibles, scissor-cutting may thus require at least as much energy per cut area as knife-cutting during cut propagation. But what about cut initiation?

Our experiments demonstrated that knife-cuts can be difficult if not impossible to initiate, because thin sheets have a low out-of-plane and in-plane stiffness, and thus easily bend and buckle before nucleating cracks; buckling loads generally increase with the flexural rigidity, *D* ∝ *t*^3^ [43]. Leaf-cutter ants may thus prefer scissor-cuts, because the mandibles then locally ‘clamp’ the sheet, so avoiding this problem altogether. Even where knife-cuts remain possible, sheet bending will incur additional elastic energy penalties, increase the cut initiation force required to meet the stress demands for crack nucleation, and thus enlarge the total energetic costs associated with cut initiation. These adverse effects can be mitigated by introducing notches into the sheet edge [either artificially, or via a preceding scissor-cut, 50– 53]; notches can increase the out-of-plane stiffness by almost a factor of six, and the in-plane stiffness by a factor of two (see figure 4b and electronic supplementary materials figure S2).

The results from *ex-vivo* cut initiation experiments and *in-silico* numerical simulations demonstrate that the mechanics of cut initiation are affected not only by the presence of a notch, but also by its angle. The narrower the angle, the larger the out-of-plane and in-plane stiffness (see figure 4 and electronic supplementary material figure S2), which increases the probability of cut initiation, and minimises the relative force increase (figure 2b,c). The notch angles at which 50 % of cuts were initiated (*P*(*Y*) = 0.5) were between 90° and 180°. This transition occurred at larger angles than the transition between *in-vivo* cutting strategies (*P*(*Y*) = 0.5 at 30−60°, figure 2a), indicating that some ants may have preferred scissor-cuts over knife-cuts, even when knife-cuts could have been initiated. This sustained preference for scissor-cuts may thus be driven by the need to initiate cuts with low forces more so than by the ability to initiate cuts at all.

## 5 Conclusion and outlook

Leaf-cutter ants use different cutting strategies to forage on thin leaves. Cuts into leaves with wide notches are typically initiated using scissor-cutting, before the ant positions itself on either side of the leaf lamina and propagates the cut via knife-cutting [49, 50]. Leaves with narrow notches, in turn, are knife-cut right away. This behavioural difference can be rationalised by considering the mechanics of thin-sheet cutting: knife-cutting thin sheets with wide notches requires high initiation forces— if cuts are initiated at all. The effects of an increased initiation force can be substantial: smaller ants would no longer be able to initiate cuts into two-thirds of the plant species they are otherwise able to cut, and this substantial negative effect provides a plausible explanation for the strong preference for scissor-cuts. We thus posit that scissor-cutting may have partially evolved as a strategy to avoid large out-of-plane deformations when initiating cuts into thin leaf laminae and flower petals. There are other potential behaviours that could reduce the risk of buckling, such as leaf edge support using the limbs, or a variation in mandible kinematics during cutting. These behaviours may be explored in work.

## Acknowledgments

This study is part of a project that has received funding from the European Research Council (ERC) under the European Union’s Horizon 2020 Research and Innovation Programme (grant agreement no. 851705) to D.L., and a Human Frontier Science Programme Young Investigator Award (grant no. RGY0073/2020) to M.B. and D.L.

## References

[1] Hung KLJ, Kingston JM, Albrecht M, Holway DA, Kohn JR. 2018 The worldwide importance of honey bees as pollinators in natural habitats. Proceedings of the Royal Society B: Biological Sciences 285: 20172140.

[2] Weisser W, Siemann E. 2004 The various effects of insects on ecosystem functioning. In: Insects and ecosystem function, Springer. pp. 3–24.

[3] Yang LH, Gratton C. 2014 Insects as drivers of ecosystem processes. Current Opinion in Insect Science 2: 26–32.

[4] Oerke EC. 2006 Crop losses to pests. Journal of Agricultural Science 144: 31–43.

[5] Dhaliwal G, Jindal V, Dhawan A. 2010 Insect pest problems and crop losses: changing trends. Indian Journal of Ecology 37: 1–7.

[6] Sharma S, Kooner R, Arora R. 2017 Insect pests and crop losses. In: Breeding insect resistant crops for sustainable agriculture, Springer. pp. 45–66.

[7] Oliveira CM, Auad AM, Mendes SM, Frizzas MR. 2014 Crop losses and the economic impact of insect pests on brazilian agriculture. Crop protection 56: 50–54.

[8] Fürstenberg-Hägg J, Zagrobelny M, Bak S. 2013 Plant defense against insect herbivores. International Journal of Molecular Sciences 14: 10242–10297.

[9] Wiens JJ, Lapoint RT, Whiteman NK. 2015 Herbivory increases diversification across insect clades. Nature Communications 6: 1–7.

[10] Bernays EA. 1991 Evolution of insect morphology in relation to plants. Philosophical Transactions of the Royal Society of London. Series B: Biological Sciences 333: 257–264.

[11] Federle W, Rohrseitz K, Hölldobler B. 2000 Attachment forces of ants measured with a centrifucge: better ‘wax-runners’ have a poorer attachment to a smooth surface. The Journal of Experimental Biology 203: 505–512.

[12] War AR, Paulraj MG, Ahmad T, Buhroo AA, Hussain B, Ignacimuthu S, Sharma HC. 2012 Mechanisms of plant defense against insect herbivores. Plant Signaling & Behavior 7: 1306–1320.

[13] Hillerton JE, Reynolds S, Vincent JF. 1982 On the indentation hardness of insect cuticle. Journal of Experimental Biology 96: 38–45.

[14] Cribb BW, Stewart A, Huang H, Truss R, Noller B, Rasch R, Zalucki MP. 2008 Insect mandibles–comparative mechanical properties and links with metal incorporation. Naturwissenschaften 95: 17–23.

[15] Schofield R, Bailey J, Coon JJ, Devaraj A, Garrett RW, Goggans MS, Hebner MG, Lee BS, Lee D, Lovern N, et al. 2021 The homogenous alternative to biomineralization: Zn-and mn-rich materials enable sharp organismal “tools” that reduce force requirements. Scientific Reports 11: 1–23.

[16] Ehrlich PR, Raven PH. 1964 Butterflies and plants: a study in coevolution. Evolution 18: 568–608.

[17] Futuyma DJ, Agrawal AA. 2009 Macroevolution and the biological diversity of plants and herbivores. Proceedings of the National Academy of Sciences of the U.S.A 106: 18054–18061.

[18] Greene E. 1989 A diet-induced developmental polymorphism in a caterpillar. Science 243: 643–646.

[19] Thompson DB. 1999 Genotype–environment interaction and the ontogeny of diet-induced phenotypic plasticity in size and shape of melanoplus femurrubrum (orthoptera: Acrididae). Journal of Evolutionary Biology 12: 38–48.

[20] Hochuli DF. 2001 Insect herbivory and ontogeny: how do growth and development influence feeding behaviour, morphology and host use? Austral Ecology 26: 563–570.

[21] Agarwala B. 2007 Phenotypic plasticity in aphids (homoptera: Insecta): Components of variation and. Current science 93.

[22] Thompson DB. 2019 Diet-induced plasticity of linear static allometry is not so simple for grasshoppers: genotype–environment interaction in ontogeny is masked by convergent growth. Integrative and Comparative Biology 59: 1382–1398.

[23] Bernays E. 1986 Diet-induced head allometry among foliage-chewing insects and its importance for graminivores. Science 231: 495–497.

[24] Bernays E, Hamai J. 1987 Head size and shape in relation to grass feeding in acridoidea (orthoptera). International Journal of Insect Morphology and Embryology 16: 323–330.

[25] Thompson DB. 1992 Consumption rates and the evolution of diet-induced plasticity in the head morphology of melanoplus femurrubrum (orthoptera: Acrididae). Oecologia 89: 204–213.

[26] Prasannakumar I, Kodandaramaiah U. 2024 Adaptive phenotypic plasticity of mandibles with respect to host plants. Arthropod-Plant Interactions 18: 77–88.

[27] Allen PE, Cui Q, Miller CW. 2021 Evidence of a rapid and adaptive response of hemipteran mouthparts to a physical barrier. Journal of Evolutionary Biology 34: 653–660.

[28] Costa AN, Vasconcelos HL, Vieira-Neto EH, Bruna EM. 2008 Do herbivores exert top-down effects in neotropical savannas? estimates of biomass consumption by leaf-cutter ants. Journal of vegetation science 19: 849–854.

[29] Wilson EO. 1980 Caste and division of labor in leaf-cutter ants (Hymenoptera: Formicidae: Atta): I. the overall pattern in A. Sextens. Behavioral Ecology and Sociobiology 7: 143–156.

[30] Wirth R, Herz H, Ryel RJ, Beyschlag W, Hölldobler B. 2003 Herbivory of Leaf-Cutting Ants: A Case Study on Atta colombica in the Tropical Rainforest of Panama. Berlin, Heidelberg, New York: Springer Science & Business Media.

[31] Hölldobler B, Wilson EO. 2010 The leafcutter ants: civilization by instinct. W. W. Norton & Company, New York City, NY, USA.

[32] Nichols-Orians CM, Schultz JC. 1989 Leaf toughness affects leaf harvesting by the leaf cutter ant, Atta cephalotes (l.) (Hymenoptera: Formicidae). Biotropica 21: 80–83.

[33] Clark E. 2006 Dynamic matching of forager size to resources in the continuously polymorphic leaf-cutter ant, Atta colombica (hymenoptera, formicidae). Ecological Entomology 31: 629–635.

[34] Roces F, Núñez JA. 1993 Information about food quality influences load-size selection in recruited leaf-cutting ants. Animal Behaviour 45: 135–143.

[35] Roces F. 1990 Leaf-cutting ants cut fragment sizes in relation to the distance from the nest. Animal Behaviour 40: 1181–1183.

[36] Lewis OT, Martin M, Czaczkes TJ. 2008 Effects of trail gradient on leaf tissue transport and load size selection in leaf-cutter ants. Behavioural Ecology 19: 805–809.

[37] Dussutour A, Deneubourg JL, Beshers S, Fourcassié V. 2009 Individual and collective problem-solving in a foraging context in the leaf-cutting ant Atta colombica. Animal cognition 12: 21–30.

[38] Moll K, Roces F, Federle W. 2010 Foraging grass-cutting ants (Atta vollenweideri) maintain stability by balancing their loads with controlled head movements. Journal of Comparative Physiology A 196: 471–480.

[39] Moll K, Roces F, Federle W. 2013 How load-carrying ants avoid falling over: mechanical stability during foraging in Atta vollenweideri grass-cutting ants. Public Library of Science One 8: e52816.

[40] Römer D, Exl R, Roces F. 2023 Two feedback mechanisms involved in the control of leaf fragment size in leaf-cutting ants. Journal of Experimental Biology 226: jeb244246.

[41] Onoda Y, Westoby M, Adler PB, Choong AMF, Clissold FJ, Cornelissen JHC, Díaz S, Dominy NJ, Elgart A, Enrico L, Fine PVA, Howard JJ, Jalili A, Kitajima K, Kurokawa H, McArthur C, Lucas PW, Markesteijn L, Pérez-Harguindeguy N, Poorter L, Richards L, Santiago LS, Sosinski Jr EE, Van Bael SA, Warton DI, Wright IJ, Joseph Wright S, Yamashita N. 2011 Global patterns of leaf mechanical properties. Ecology Letters 14: 301–312.

[42] Hua L, He P, Goldstein G, Liu H, Yin D, Zhu S, Ye Q. 2020 Linking vein properties to leaf biomechanics across 58 woody species from a subtropical forest. Plant Biology 22: 212–220.

[43] Bhaskar K, Varadan T. 2021 Plates: theories and applications. Springer Nature.

[44] Chapman R. 1995 Mechanics of food handling by chewing insects. In: Chapman RF, de Boer G, editors, Regulatory mechanisms in insect feeding, Springer. pp. 3–31.

[45] Blanke A, Machida R, Szucsich NU, Wilde F, Misof B. 2015 Mandibles with two joints evolved much earlier in the history of insects: dicondyly is a synapomorphy of bristletails, silverfish and winged insects. Systematic Entomology 40: 357–364.

[46] Krenn HW. 2019 Insect mouthparts: form, function, development and performance. Springer Nature.

[47] Kang V, Püffel F, Labonte D. 2023 Three-dimensional kinematics of leaf-cutter ant mandibles: not all dicondylic joints are simple hinges. Philosophical Transactions of the Royal Society B 378: 20220546.

[48] Sanson G. 2006 The biomechanics of browsing and grazing. American Journal of Botany 93: 1531–1545.

[49] Tautz J, Roces F, Hölldobler B. 1995 Use of a sound-based vibratome by leaf-cutting ants. Science 267: 84.

[50] Schofield RM, Emmett KD, Niedbala JC, Nesson M. 2011 Leaf-cutter ants with worn mandibles cut half as fast, spend twice the energy, and tend to carry instead of cut. Behavioral Ecology and Sociobiology 65: 969–982.

[51] Lake GJ, Yeoh OH. 1978 Measurement of rubber cutting resistance in the absence of friction. International Journal of Fracture 14: 509–526.

[52] Zhang B, Shiang CS, Yang SJ, Hutchens SB. 2019 Y-shaped cutting for the systematic characterization of cutting and tearing. Experimental Mechanics 59: 517–529.

[53] Püffel F, Walthaus OK, Kang V, Labonte D. 2023 Biomechanics of cutting: sharpness, wear sensitivity, and the scaling of cutting forces in leaf-cutter ant mandibles. Philosophical Transactions of the Royal Society B 378: 20220547.

[54] Eidmann H. 1936 Zur kenntnis der blattschneiderameise atta sexdens l., insbesondere ihrer ökologie. Zeitschrift für angewandte Entomologie 22: 385–436.

[55] Barrer P, Cherrett JM. 1972 Some factors affecting the site and pattern of leaf-cutting activity in the ant Atta cephalotes l. Physiological Entomology 47: 15–27.

[56] Püffel F, Pouget A, Liu X, Zuber M, van de Kamp T, Roces F, Labonte D. 2021 Morphological determinants of bite force capacity in insects: a biomechanical analysis of polymorphic leaf-cutter ants. Journal of the Royal Society Interface 18: 20210424.

[57] Plum F, Labonte D. 2021 scant—an open-source platform for the creation of 3d models of arthropods (and other small objects). PeerJ 9: e11155.

[58] Püffel F, Roces F, Labonte D. 2023 Strong positive allometry of bite force in leaf-cutter ants increases the range of cuttable plant tissues. Journal of Experimental Biology 226: jeb245140.

[59] Brown RHJ. 1963 Jumping arthopods. Times Science Review : 6–7.

[60] Roces F, Hölldobler B. 1994 Leaf density and a trade-off between load-size selection and recruitment behaviour in the ant Atta cephalotes. Oecologia 97: 1–8.

[61] McCarthy CT, Hussey M, Gilchrist MD. 2007 On the sharpness of straight edge blades in cutting soft solids: Part i - indentation experiments. Engineering Fracture Mechanics 74: 2205–2224.

[62] Spagnoli A, Brighenti R, Terzano M, Artoni F. 2019 Cutting resistance of soft materials: Effects of blade inclination and friction. Theoretical and Applied Fracture Mechanics 101: 200–206.

[63] Duncan TT, Sarapas JM, Defante AP, Beers KL, Chan EP. 2020 Cutting to measure the elasticity and fracture of soft gels. Soft Matter 16: 8826–8831.

[64] Williams JG, Patel Y. 2016 Fundamentals of cutting. Interface Focus 6: 20150108.

[65] Reyssat E, Tallinen T, Le Merrer M, Mahadevan L. 2012 Slicing softly with shear. Physical Review Letters 109: 244301.

[66] Goda BA, Labonte D, Bacca M. 2024 Making the cut: end effects and the benefits of slicing. Extreme Mechanics Letters 72: 102221.

[67] R Core Team. 2021 R: A Language and Environment for Statistical Computing. R Foundation for Statistical Computing, Vienna, Austria. URL https://www.R-project.org/.

[68] Field A, Miles J, Field Z. 2012 Discovering Statistics Using R. SAGE Publications Ltd.

[69] Sharpe WN, editor 2008 Springer Handbook of Experimental Solid Mechanics. Springer Science & Business Media, 1 edition.

[70] Clissold F. 2007 The biomechanics of chewing and plant fracture: Mechanisms and implications. Advances in Insect Physiology 34.

[71] Peeters PJ, Sanson G, Read J. 2007 Leaf biomechanical properties and the densities of herbivorous insect guilds. Functional Ecology : 246–255.

[72] Gronenberg W, Paul J, Just S, Hölldobler B. 1997 Mandible muscle fibers in ants: fast or powerful? Cell & Tissue Research 289: 347–361.

[73] Paul J, Gronenberg W. 1999 Optimizing force and velocity: Mandible muscle fibre attachments in ants. The Journal of Experimental Biology 202: 797–808.

[74] Garrett RW, Carlson KA, Goggans MS, Nesson MH, Shepard CA, Schofield RM. 2016 Leaf processing behaviour in Atta leaf-cutter ants: 90% of leaf cutting takes place inside the nest, and ants select pieces that require less cutting. Royal Society Open Science 3: 1–12.

[75] Hart AG, Ratnieks FL. 2001 Task partitioning, division of labour and nest compartmentalisation collectively isolate hazardous waste in the leafcutting ant Atta cephalotes. Behavioral Ecology and Sociobiology 49: 387–392.

[76] Camargo R, Forti LC, Lopes J, Andrade A, Ottati A. 2007 Age polyethism in the leaf-cutting ant Acromyrmex subterraneus brunneus forel, 1911 (hym., formicidae). Journal of Applied Entomology 131: 139–145.

[77] Doran C, McCormack B, Macey A. 2004 A simplified model to determine the contribution of strain energy in the failure process of thin biological membranes during cutting. Strain 40: 173–179.

[78] Atkins AG, Xu X, Jeronimidis G. 2004 Cutting, by ‘pressing and slicing,’of thin floppy slices of materials illustrated by experiments on cheddar cheese and salami. Journal of Materials Science 39: 2761–2766.

[79] Atkins A, Xu X. 2005 Slicing of soft flexible solids with industrial applications. International Journal of Mechanical Sciences 47: 479–492.

[80] Atkins AG, Mai YW. 1979 On the guillotining of materials. Journal of Materials Science 14: 2747–2754.

[81] Lucas P, Pereira B. 1990 Estimation of the fracture toughness of leaves. Functional Ecology 4: 819–822.

