## Supplementary Materials for "Behavioural biomechanics: leaf-cutter ant cutting behaviour depends on leaf edge geometry"

### Supplementary material

#### Sensor calibration & drift correction

Prior to all calibrations and experiments, the fibre-optic displacement sensor was positioned approximately 400 $\mu$m above the unloaded bending beam to ensure consistent sensor readings (thickness: 0.5 mm, width: 10.2 mm, free length: 25.3 mm). The sensor was then calibrated with ten calibration weights, ranging from 10 to 392 mN (1-40 g, Kern & Sohn, Balingen, Germany), exceeding the range of cut initiation and steady-state forces (20-385 mN). The weights were suspended from a polymer mount located at the free end of the bending beam, and the average displacement was extracted from the force recording across a duration of 2 s. Sensor drift was accounted for by implementing a linear drift correction based on the sensor output for the unloaded beam, at the beginning and end of the recording. Both linear and quadratic regression models were tested to characterise the relationship between sensor distance and calibration force. The model which yielded the lower Akaike information criterion was then selected for calibration; in ten out of eleven cases, the quadratic model was preferred. All selected regression models yielded an R-squared of at least 0.998, with an approximate slope, or sensor 'compliance', of 0.40 mm/N.

#### Steady-state cutting force measurements

Steady-state cutting force measurements were conducted similar to the cut initiation force measurements, using the same experimental setup, bending beam, and motor speed, but with three experimental differences: First, the PDMS sheets were of an intermediate thickness (207±3 $\mu$m), with only a small notch in the centre (30° and 1.5 mm deep). Second, the sheets were clamped onto a different polymer mount without a lower clamping bar and a narrower cutting region (2 mm wide and 8 mm long, see [1] figure 1c for a similar design). Third, the motor was moved for a total of 5 mm (instead of 2.5 mm) so that steady-state cutting forces could be reliably extracted as averages across 2 mm cut length. Steady-state forces ranged between 25-121 mN (figure 3 in the main text) — at the lower end comparable to a theoretical minimum predicted from the sheet thickness and fracture toughness of PDMS, 32 ± 6 mN. We note that this fracture toughness estimate, extracted from pure shear tests, may include an energetic 'overhead' associated with the creation of a larger fracture process zone than present during cutting [2]. This 'overhead' may explain why some cutting force measurements were even lower than $F_{min} = G_c t$. At the upper end, the highest steady-state cutting forces exceeded $F_{min}$ by a factor of about four; mandibles that cut with high steady-state forces also initiated significantly fewer cuts across the eight test conditions (Kendall's $\tau = -0.76$, p < 0.0001, n = 18).

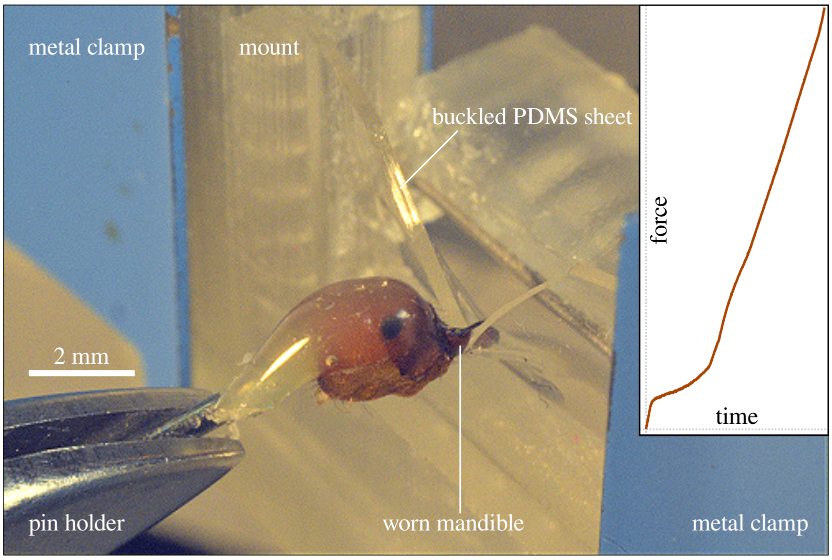


Figure 1 | *Ex-vivo* cut initiation experiments that simulated knife-cutting were performed, using leaf-cutter ant mandibles as cutting tool and PDMS pseudoleaves with varying sheet edge geometry as cutting substrate. For pseudoleaves with wide notches or mandibles with high degrees of mandibular wear, cuts often failed to initiate, and the pseudoleaves buckled substantially out-of-plane instead. Sheet buckling was associated with a visible drop in 'stiffness' in the force-time graphs.

### References

[1] Püffel F, Walthaus OK, Kang V, Labonte D. 2023 Biomechanics of cutting: sharpness, wear sensitivity, and the scaling of cutting forces in leaf-cutter ant mandibles. *Philosophical Transactions of the Royal Society B* **378**: 20220547.

[2] Goda BA, Ma Z, Fregonese S, Bacca M. 2024 Cutting soft matter: Scaling relations controlled by toughness, friction and wear. *Soft Matter* : DOI: 10.1039/D4SM00279B.

[3] Field A, Miles J, Field Z. 2012 Discovering Statistics Using R. SAGE Publications Ltd.

Table 1 | Results of binary logistic regression analysis of form $P(Y) = 1/(1+e^{-(b_0+b_1X_1+...+b_nX_n)})$, describing the probability of 'knife-cut' initiation during *in-vivo* cutting-behavioural assays and successful cut initiations during *ex-vivo* cutting force experiments. Significance levels of the regression coefficients, $b_i$, are denoted by an asterisk (. p < 0.05, * p < 0.01, ** p < 0.001, *** p < 0.0001), calculated based on the $z$-statistic. The 'odds ratio' reflects the change of odds, $P(Y)/P(\text{No}\,Y)$, associated with a unit increase of $X_i$. Values greater than unity indicate an increase in odds; conversely, values smaller than unity indicate a decrease in odds (95 % confidence intervals are provided in parentheses). Whether or not the successive addition of model predictors ($X_1$ = notch angle in degrees, $X_2$ = sheet thickness in micrometers) led to a significant improvement of the model was assessed via $\chi^2$-tests following Fields et al. [3].

<table>
<tr><th>Experiment</th><th>Substrate</th><th>Event $Y$ *vs* No event $Y$</th><th>Predictor $X_i$</th><th>Coefficient $b_i$ (standard error)</th><th>Odds ratio (95% CI)</th><th>Model comparison</th><th>$\chi^2$-statistic</th><th>P value</th></tr>
<tr><td>Cutting trial</td><td>PDMS</td><td>'Knife' *vs* 'Scissors'</td><td>(-)<br>Notch angle<br>Sheet thickness</td><td>$b_0$ = 2.3958*** (0.4316)<br>$b_1$ = -0.0301*** (0.0030)<br>$b_2$ = -0.0040** (0.0015)</td><td>(-)<br>0.970 (0.964|0.976)<br>0.996 (0.993|0.999)</td><td>$f(X_1,X_2)$ *vs* $f(X_1)$</td><td>$\chi^2_1$ = 6.87</td><td><0.01</td></tr>
<tr><td>Cutting trial</td><td>Laurel</td><td>'Knife' *vs* 'Scissors'</td><td>(-)<br>Notch angle</td><td>$b_0$ = 1.0838*** (0.2781)<br>$b_1$ = -0.0277*** (0.0041)</td><td>(-)<br>0.973 (0.964|0.980)</td><td>$f(X_1)$ *vs* intercept</td><td>$\chi^2_1$ = 75.95</td><td><0.001</td></tr>
<tr><td>Force experiment</td><td>PDMS</td><td>Cut *vs* No cut</td><td>(-)<br>Notch angle<br>Sheet thickness</td><td>$b_0$ = 1.7948 (1.3350)<br>$b_1$ = -0.0515*** (0.0129)<br>$b_2$ = 0.0214** (0.0067)</td><td>(-)<br>0.950 (0.918|0.969)<br>1.022 (1.010|1.038)</td><td>$f(X_1,X_2)$ *vs* $f(X_1)$</td><td>$\chi^2_1$ = 18.19</td><td><0.001</td></tr>
</table>

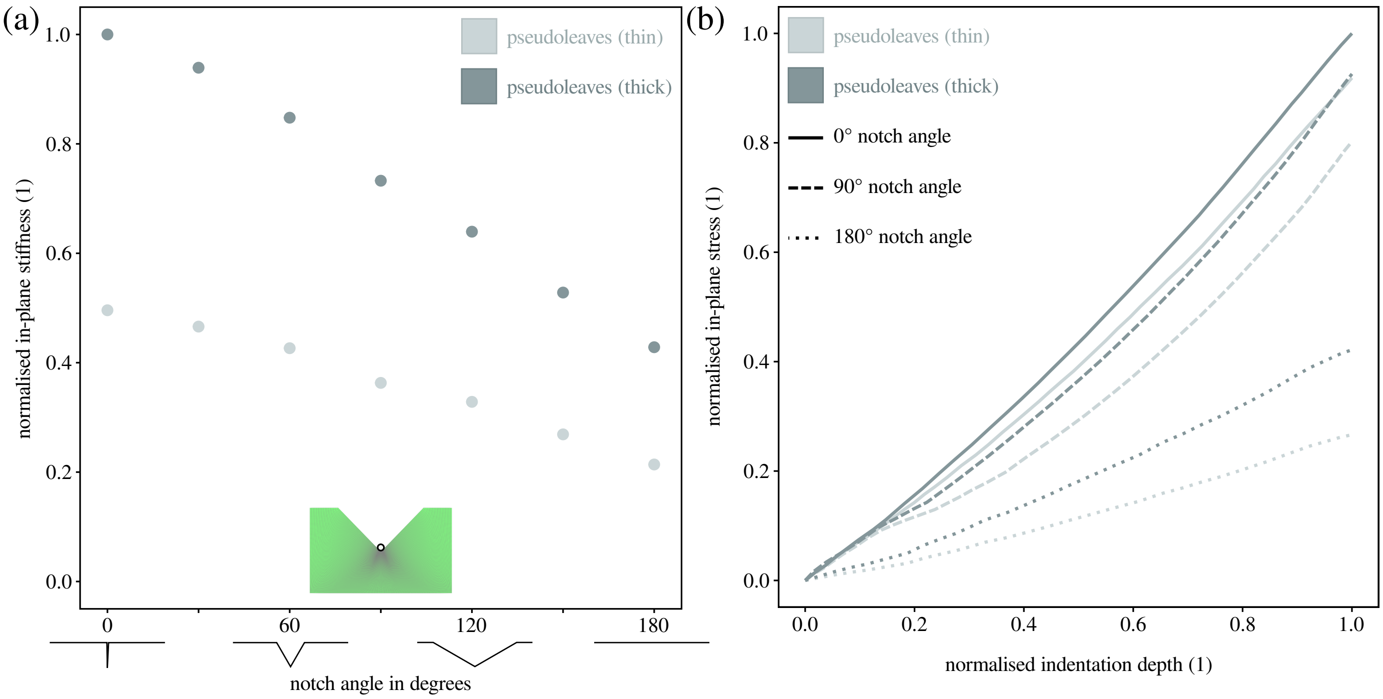


Figure 2 | To better understand the effects of notch angle and sheet thickness on the mechanics of cut initiation, we performed Finite Element Analysis on the rectangular sheet geometries used during *ex-vivo* cutting experiments. *(a)* Upon displacing the notch centres by a fixed distance of 200 $\mu$m in the direction of cutting, the resulting reaction force per unit displacement, the in-plane stiffness, was maximum for thick sheets with narrow notches, and monotonously decreased with increasing notch angle; stiffness values were normalised with the global maximum. In-plane stiffness approximately halved between thick and thin sheets and between 0° and 180° notch angles. *(b)* As a consequence, the maximum tensile stresses, which generally increased with indentation depth, were substantially larger for narrower angles, suggesting that sharp notches not only reduce out-of-plane deformations but also facilitate cut initiation by concentrating stresses around the forming crack; stress values were normalised with the global maximum and indentation depth with the total displacement of 200 $\mu$m.
